# AlphaPulldown2 – A General Pipeline for High-Throughput Structural Modeling

**DOI:** 10.1101/2024.11.28.625873

**Authors:** Dmitry Molodenskiy, Valentin J. Maurer, Dingquan Yu, Grzegorz Chojnowski, Stefan Bienert, Gerardo Tauriello, Konstantin Gilep, Torsten Schwede, Jan Kosinski

**Author notes:** These authors contributed equally to this work (Dmitry Molodenskiy, Valentin J. Maurer, Dingquan Yu).

## Abstract

AlphaPulldown2 streamlines protein structural modeling by automating workflows, improving code adaptability, and optimizing data management for large-scale applications. It introduces an automated Snakemake pipeline, compressed data storage, support for additional modeling backends like UniFold and AlphaLink2, and a range of other improvements. These upgrades make AlphaPulldown2 a versatile platform for predicting both binary interactions and complex multi-unit assemblies.

**Availability:** AlphaPulldown2 is freely available at https://github.com/KosinskiLab/AlphaPulldown.

## 1. Introduction

Recent advancements in Artificial Intelligence (AI)-based structural prediction, driven by tools such as AlphaFold2 (Jumper et al., 2021), RoseTTAFold (Baek et al., 2021), and ColabFold (Mirdita et al., 2022), have remarkably improved our capacity to predict protein– protein interactions (PPIs) and the architecture of protein complexes. The associated confidence scores can also be used to predict whether two proteins would interact, facilitating high-throughput computational screens of PPIs (Humphreys et al., 2021; Burke et al., 2023; Yu et al., 2023; Razew et al., 2024).

We previously introduced the AlphaPulldown Python package (Yu et al., 2023) to streamline PPI screens and facilitate high-throughput modeling of higher-order complexes using AlphaFold-Multimer (Evans et al., 2021). AlphaPulldown separates the AlphaFold pipeline into CPU-based calculation of input features (multiple sequence alignments (MSAs) and templates) and GPU-based structure prediction, reducing computational time. It offers four modes: pulldown, all-versus-all, homo-oligomer, and custom (Supplementary Fig. 1). The pulldown mode screens interactions between one or more “bait” proteins and a list of candidates, mimicking pulldown assays. The all-versus-all mode automatically predicts pairwise PPIs between all proteins in a provided list, useful for interaction network prediction. The homo-oligomer mode facilitates the modeling of alternative oligomeric states. The custom mode allows flexible input combinations of proteins or fragments. AlphaPulldown reuses input features pre-calculated for full-length proteins when modeling fragments, avoiding costly recalculation, and retains original residue numbering in the resulting models. It includes an integrated analysis pipeline that enriches native AlphaFold scores with additional evaluation metrics such as pDockQ (Bryant et al., 2022) and physical parameters of interfaces (Malhotra et al., 2021; Agirre et al., 2023), generating a graphical summary in a Jupyter notebook for comprehensive analysis of model confidence and interaction properties. AlphaPulldown has been utilized in a range of applications, demonstrating its effectiveness in PPI screens, modeling individual complexes, and interface scoring (Kadhim et al., 2024; Bonchuk et al., 2024; Sellés-Baiget et al., 2024; Lapcik et al., 2024; Seidel et al., 2023).

Despite its utility, challenges in automation, code adaptability, and management of the resulting models and data persisted. Moreover, new customized modeling protocols have emerged for adjusting modeling outcomes using modified MSAs, templates, or distance restraints (Stahl et al., 2024; Mirabello et al., 2024). In response, we introduce AlphaPulldown version 2.0, incorporating significant improvements in user experience, modeling capabilities, and available modeling and evaluation features. This updated version offers a comprehensive suite for modeling both binary PPIs and multi-unit assemblies, positioning it as a comprehensive toolbox for high-throughput protein structure prediction.

## 2. Software Description

### 2.1. Usability Improvements and Software Management

#### 2.1.1. Automation and New Configuration Syntax

In large-scale structural modeling, managing multiple modeling tasks (jobs) and computational resources becomes a bottleneck. Thus, to further automate AI-based structural modeling, we developed an automated scalable and reproducible pipeline for AlphaPulldown2 using the Snakemake workflow management system (Mölder et al., 2021). This pipeline replicates the original AlphaPulldown workflow but now runs all steps automatically based on an initial configuration (Fig. 1). Snakemake internally uses Singularity containers (Kurtzer et al., 2017) to ensure reproducibility and compatibility with various compute architectures, including cloud or local clusters. It reschedules jobs with settings adjusted based on failure reasons and resumes from saved checkpoints. Snakemake also simplifies the installation process, as AlphaPulldown is now available on DockerHub (Merkel, 2014) and is automatically installed during the first execution.

**Fig. 1.**
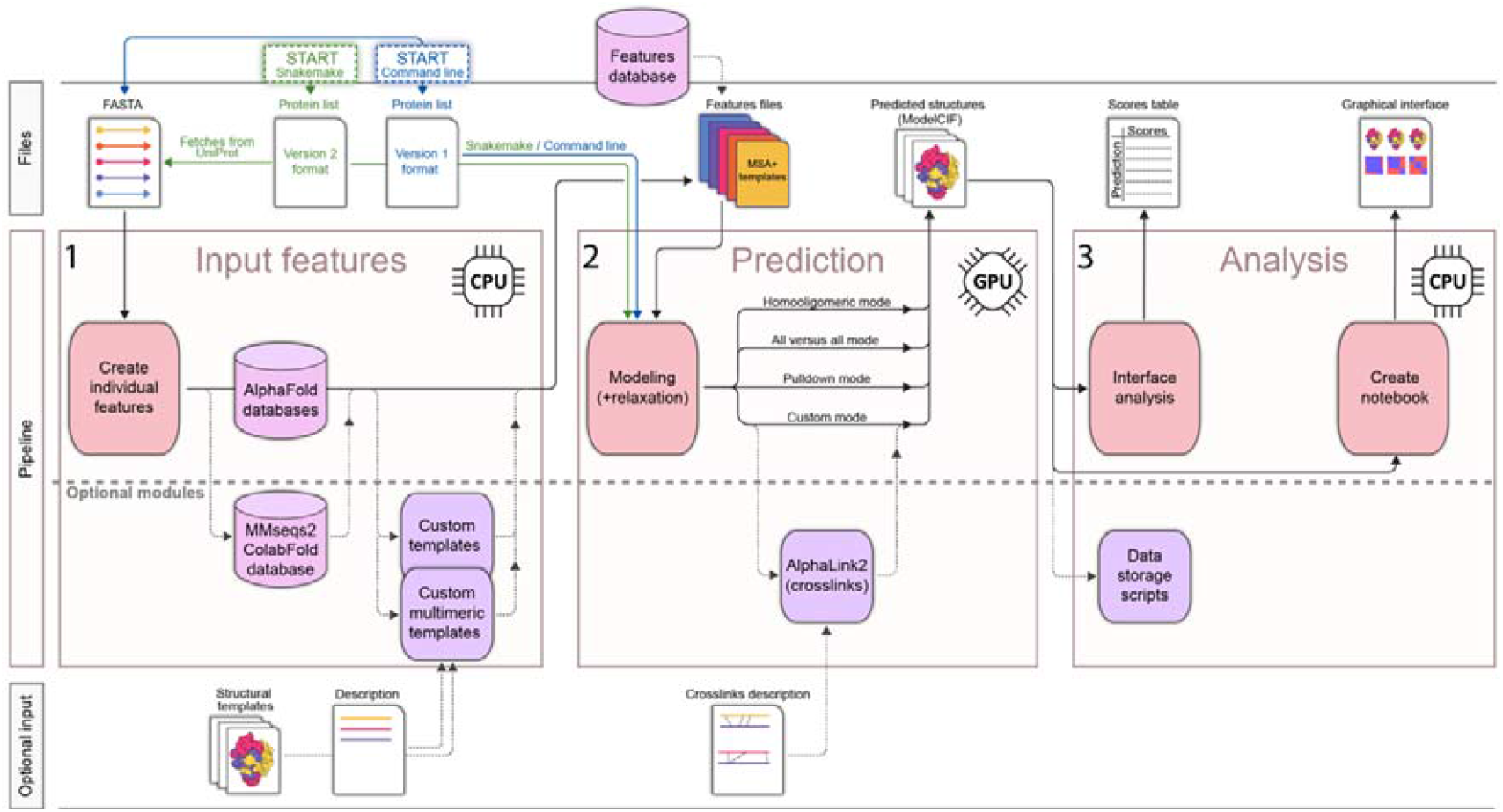
The workflow of structural modeling with AlphaPulldown2. For better allocation of resources, AlphaPulldown2 separates the AlphaFold pipeline into CPU-based calculation of input features and results analysis (1,3) and GPU-based structure prediction (2). The pipeline can use a new Version 2 input format that can define all Version 1 modes using simplified syntax. Additional input like cross-links and multimeric templates can be provided. The resulting models in ModelCIF format and confidence scores are summarized using an interactive, graphical interface. Features and output files are compressed to decrease storage requirements. The entire pipeline can be run automatically using Snakemake.

We also introduced a new syntax for configuring the modeling jobs, which unifies all AlphaPulldown modes—pulldown, homo-oligomer, all-versus-all, and custom—into a single, flexible format (Supplementary Fig. 1). Within this format, AlphaPulldown2 additionally allows users to provide UniProt IDs of the input proteins (The UniProt Consortium, 2023), and the sequences will be fetched automatically. This simplifies input preparation and allows users to define complex modeling scenarios with ease. While the original scripts and syntax remain available, the Snakemake pipeline and new configuration syntax enhance automation and usability.

#### 2.1.2. Storage Management

High-throughput AI-based structure prediction requires extensive storage, as input features and output files with confidence scores can easily accumulate to terabytes of data. To mitigate this, we implemented several space-saving measures. Input features in PKL format (Python object serialization) are now compressed using the XZ format with the LZMA2 compression algorithm, reducing file sizes by 97.7% (55⍰GB instead of 2.4⍰TB for 20,581 human proteins). Output PKL files are compressed too, and usually unnecessary data (aligned confidence probabilities, distograms, and masked MSAs) are removed before saving. This reduces the average size of output directories by 97%. These compressions are optional, with a script available for cleaning and compressing the data later. For instance, in the case of modeling 1,000 random protein pairs, storage requirements dropped from 3.5⍰TB to 102⍰GB with AlphaPulldown2.

#### 2.1.3. Improved Code Organization and Testing Framework

To support community interest, feature requests, and our development goals, we have restructured the AlphaPulldown codebase. AlphaPulldown was initially built on AlphaFold-Multimer, but other implementations like OpenFold (Ahdritz et al., 2024) and UniFold (Li et al., 2022) have since emerged. UniFold, in particular, offers the capabilities of AlphaFold and includes code for training and fine-tuning AlphaFold weights. New tools such as AlphaFold3 (Abramson et al., 2024), OpenFold Multimer, and HelixFold3 (Liu et al., 2024) are also becoming available. To ensure that AlphaPulldown is able to timely incorporate these folding programs in the future, we reorganized the architecture of AlphaPulldown2 to allow flexible integration of various modeling backends. Input features can be generated through a unified pipeline using AlphaFold2 or MMSeqs2 (Steinegger and Söding, 2017; Mirdita et al., 2022) and passed to the user’s chosen backend. For example, we added backend support for UniFold and AlphaLink2 (see section 2.3), and additional backends can be integrated using this modular system.

Further improvements include code refactoring for clarity and efficiency, the removal of redundancies, and the addition of automated tests. We also implemented continuous integration and continuous delivery/deployment (CI/CD) pipelines to streamline ongoing development. AlphaPulldown docker images are automatically built and pushed to DockerHub upon each new software release to the AlphaPulldown GitHub repository, ensuring that the software remains up to date. Lastly, we expanded the usage manual to provide more comprehensive documentation.

### 2.2. Ensuring FAIR Principles through ModelCIF Support

In the era of ubiquitous structural modeling, one major challenge is ensuring that the resulting data are both reproducible and accessible in a standardized format. The lack of detailed metadata in the traditional PDB format limits the ability to reproduce results and assess model reliability.

To address this, we have added support for the ModelCIF format (Vallat et al., 2023). ModelCIF, an extension of the PDBx/mmCIF dictionary (Westbrook et al., 2022), is specifically designed to accommodate computationally generated models. ModelCIF aligns with the FAIR (Findable, Accessible, Interoperable, and Reusable) principles, enabling detailed documentation of the modeling process, including software versions, parameters, and information about used sequence and template databases. The global confidence scores as well as local pLDDT and pairwise alignment error matrices are stored in the files as well. This extension ensures that models predicted using AlphaPulldown2 can be stored with all necessary metadata for future replication and analysis. The introduction of ModelCIF support also ensures that AlphaPulldown2-generated models are fully compatible with repositories like ModelArchive (https://www.modelarchive.org/) (Schwede et al., 2009), facilitating model deposition. This integration not only facilitates compliance with FAIR principles but also provides a standardized and reliable way to store, share, and evaluate high-throughput structural modeling data.

### 2.3. Cross-Link-Driven Modeling through the Integration of AlphaLink2

Cross-linking mass spectrometry (XL-MS) is a powerful experimental technique used to study PPIs and macromolecular complexes (Graziadei and Rappsilber, 2022). In XL-MS, chemical cross-linkers covalently bind specific amino acid residues that are in proximity. By analyzing these cross-linked peptides via mass spectrometry, the cross-linked residue pairs can be identified and used as distance restraints in structural modeling.

AlphaLink2 (Stahl et al., 2024) is a modified version of AlphaFold2, based on the UniFold implementation, that incorporates cross-linking data as modeling restraints within the AlphaFold2 neural network. This enhancement improves the accuracy of structural models, particularly for large protein complexes and challenging PPIs.

In AlphaPulldown2, we have implemented AlphaLink2 as an optional backend, enabling seamless integration of cross-link-driven modeling into the existing workflow. The process begins with feature generation using the standard AlphaFold2 or MMSeqs2 pipelines, after which these features are passed to AlphaLink2, where cross-linking restraints are applied to guide structure prediction. This integration allows users to perform both unconstrained and restraint-driven modeling within the same software framework, offering flexibility for various use cases while benefiting from the full range of features provided by AlphaPulldown2.

### 2.4. Extended Modeling Applications

AlphaPulldown2 offers a wide range of parameters and features that expand the capabilities of AlphaFold2. These include options for adjusting the number of recycles, specifying the number of output models, and selecting between different sequence search methods, such as the standard AlphaFold2 pipeline or the faster MMSeqs2. Users can also choose between different template search methods, like the faster HMMER (Finn et al., 2015) or the more accurate HHsearch (Steinegger et al., 2019). These customizable parameters enable users to optimize modeling protocols, whether prioritizing speed or accuracy.

AlphaPulldown2 also introduces enhanced functionalities for controlling predictions. Users can adjust the MSA by toggling the pairing of sequences from the same species or modifying MSA depth to increase the diversity of models (Monteiro da Silva et al., 2024). Additionally, custom templates, including multimeric templates, can be used. The multimeric templates allow users to impose the relative orientation of template chains on their models, improving the accuracy of complexes that cannot be accurately predicted by AlphaFold2 alone (Mirabello et al., 2024). This feature is particularly useful for refining pre-calculated models of smaller complexes or partially resolved experimental structures by adding missing regions or proteins. Additionally, it can automatically remove clashing residues and/or regions of low confidence (low pLDDT scores) when previous AlphaFold2 models are provided as templates. Since the reliance of AlphaFold2 on templates varies with the depth of the MSA, AlphaPulldown2 includes an automated mode that samples a gradient of MSA depths. This mode enables users to fine-tune the degree to which templates influence the final model. Finally, the analysis pipeline has been enriched with the commonly used and requested average pLDDT and PAE scores at protein interfaces, offering a more comprehensive assessment of PPIs and model quality.

### 2.5. Repository of Input Features for Model Organisms

The generation of input MSA and template features is computationally intensive and often redundantly performed by different labs for the same proteins. This leads to unsustainable use of resources and time. To address this, we released a web-based repository (linked from the AlphaPulldown GitHub page) of the input features for the proteomes of 12 model organisms. Users can download individual input features in compressed PKL format or entire proteomes, allowing them to proceed directly to structure prediction without the need for MSA and template generation. This approach not only accelerates workflows but also reduces global computational costs and the associated energy consumption from repeatedly generating input features for the same proteins.

## 3. Conclusions, Discussion, and Future Plans

AlphaPulldown2 represents a significant step forward in high-throughput AI-based structural modeling, offering a versatile and customizable platform for protein complex predictions. With its new automation features, support for different modeling backends, and the ability to integrate experimental data like cross-links, AlphaPulldown2 further enhances modeling quality. Its optimizations in storage management and computational efficiency also make it more sustainable for large-scale projects, reducing both time and resource consumption. Although AlphaFold3 has been released, including its source code, and reproductions like HelixFold3 and Chai-1 have emerged, they require further testing before they can be confidently incorporated into workflows such as AlphaPulldown. Additionally, the more restrictive licensing terms of these new programs may present challenges, particularly for high-throughput applications and for depositing models into public databases. In terms of modeling protein complexes, AlphaFold3 only slightly improved in accuracy over AlphaFold2. Moreover, AlphaFold3 has been shown to hallucinate false secondary structures more frequently than AlphaFold2. Thus, even though AlphaPulldown relies on AlphaFold2 and AlphaFold-Multimer, it remains a robust and reliable tool for high-throughput protein complex modeling.

Looking ahead, we plan to incorporate additional modeling backends, such as OpenFold, once its multimeric modeling mode is fully supported. We are also working on expanding the range of scoring functions and will continue to evaluate and integrate permissively licensed reproductions of AlphaFold3 as they become available. Thanks to these new modeling backends, future developments will also focus on enabling the incorporation of other molecules and post-translational modifications.

## Acknowledgements

We thank AlphaPulldown users for bug reports and feature requests. We thank Stephen Royle, Sebastian Swanson, and Kotaro Tanaka for code contributions.

## Author Contributions

DM – Conceptualization, Software, Writing – review & editing

VM – Conceptualization, Software, Writing – review & editing

DY – Conceptualization, Software, Writing – review & editing

GC – Software, Writing – review & editing

SB – Software

GT – Software

KG – Software, Visualization

TS – Funding acquisition, Supervision

JK – Conceptualization, Funding acquisition, Project administration, Supervision, Software, Writing – original draft, Writing – review & editing

---

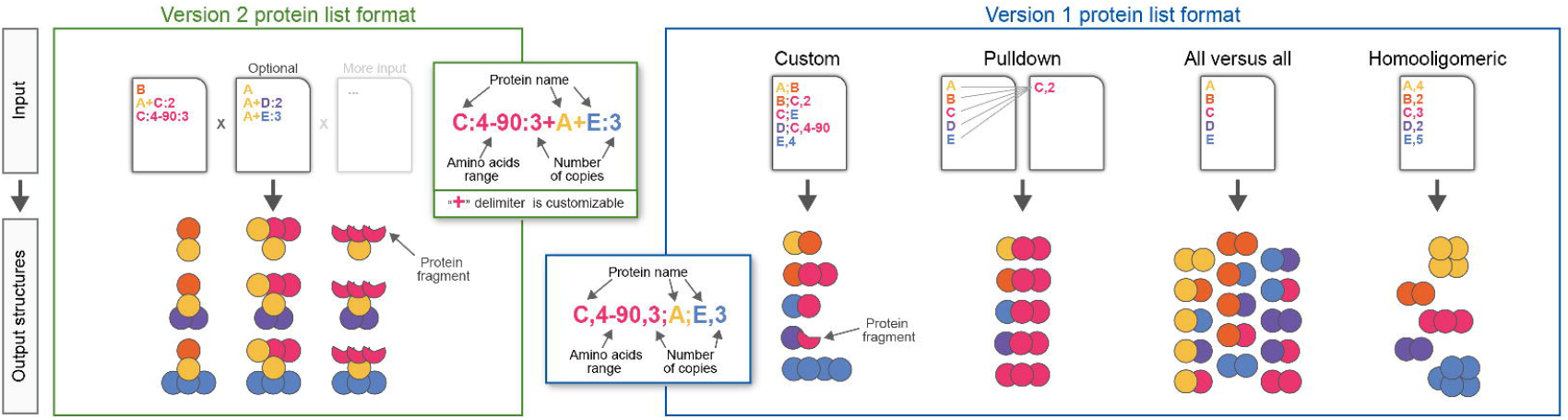

