## Supplemental Figure 1 for "AlphaPulldown2 – A General Pipeline for High-Throughput Structural Modeling"

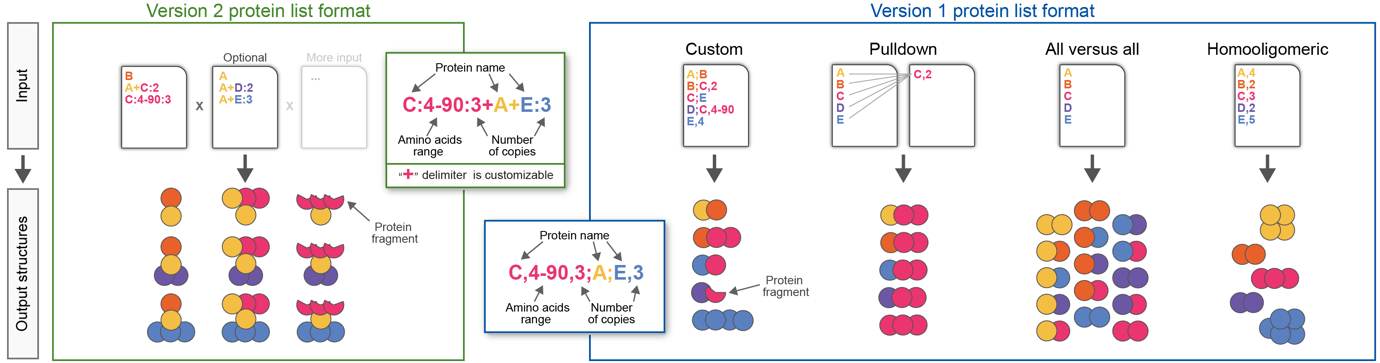

**Supplementary Fig. 1.** AlphaPulldown modeling modes and input syntax. Version 1 of AlphaPulldown (right) includes four distinct modeling modes: pulldown, all-versus-all, homo-oligomer, and custom, each requiring specific input syntax. In version 2 (left), these syntaxes are consolidated into a single, versatile format applicable to all modes. Both versions allow the selection of specific amino acid residue ranges for modeling, extracted from the full-length protein input, with the resulting models retaining the original residue numbering. AlphaPulldown2 supports both input formats to ensure backward compatibility.
